# A high-throughput exonuclease assay based on the fluorescent base analog 2-aminopurine

**DOI:** 10.1101/2022.09.12.507544

**Authors:** Margherita M. Botto, Sudarshan Murthy, Meindert H. Lamers

**Affiliations:** Cell and Chemical Biology Department, Leiden University Medical Center, Einthovenweg 20, 2333 ZC Leiden, The Netherlands

## Abstract

Exonucleases are essential enzymes that remove nucleotides from free DNA ends during DNA replication, DNA repair and telomere maintenance. Due to their essential role, they are potential targets for novel anticancer and antimicrobial drugs but have so far have been little exploited. Here we present a simple and versatile real-time exonuclease assay based on 2-aminopurine, an intrinsically fluorescent nucleotide that is quenched by neighboring bases when embedded in DNA. We show that our assay is applicable to different eukaryotic and bacterial exonucleases acting on both 3’ and 5’ DNA ends, over a wide range of protein activities and suitable for a high throughput inhibitor screening campaign. Using our assay, we discover a novel inhibitor of the *Mycobacterium tuberculosis* PHP-exonuclease that is part of the replicative DNA polymerase DnaE1. Hence, our novel assay will be a useful tool for high throughput screening for novel exonuclease inhibitors that may interfere with DNA replication or DNA maintenance.

## INTRODUCTION

Exonucleases are essential enzymes that remove mis-incorporated nucleotides during DNA replication, modify DNA during crosslink repair, mismatch repair, non-homologous end-joining, nucleotide excision repair and also during telomere maintenance (reviewed in ^1^). Exonucleases cleave the terminal nucleotide of either single or double stranded DNA and can act in either a 3’-5’ or a 5’-3’ direction. They are distinct from endonuclease that cut DNA internally. Because of their essential roles in DNA replication and DNA repair, they are potentially attractive targets for novel therapeutics that aim to interfere with DNA synthesis or DNA maintenance. However, exonucleases have thus far been little exploited, with the exception of a small number of examples, such as the exonuclease domain of the *M. tuberculosis* replicative DNA polymerase DnaE1^2^ or the exonuclease domain of the *Streptococcus pneumoniae* Pol C^3^.

To aid in the discovery of novel exonuclease inhibitors, a suitable assay is needed that can be used in a high-throughput manner, thus excluding widely used gel-based assays that are laborious and time consuming. Several real-time, fluorescence-based exonuclease assays have been reported but each of these presents different drawback that may obstruct the discovery of novel inhibitors. For example, 5′-*p*-nitrophenyl ester of TMP works well on the *E. coli* exonuclease ε^4^, but is not hydrolyzed by the exonuclease of the replicative DNA polymerase DnaE1 from *M. tuberculosis* (Lamers, unpublished results). Other real-time exonuclease assays use DNA intercalating dyes that could affect the activity of the exonuclease such as in^5^. The intercalating dye-based assay furthermore requires processive exonuclease activity, as the removal of a single nucleotide will not suffice to change the fluorescent signal. Finally, fluorescent dyes coupled to the end of the DNA substrate^6^ or fluorescent dye-quencher pairs have also been used ^7^, but here the presence of a bulky chemical group may interfere with enzymatic activity.

In this work, we present a high-throughput exonuclease assay based on 2-aminopurine (2-AP), an intrinsically fluorescent analog of adenine and guanine ^8^ (Figure 1B). The fluorescence of 2-AP is quenched by the presence of neighboring bases in the DNA ^9^, but is restored when released by an exonuclease (Figure 1A). 2-AP has been previously used to inquire nucleic acid structure ^10–13^, their dynamics ^14–16^ and enzyme activity ^17–19^, including that of exonucleases. However, thus far it has not been applied for the use of high-throughput inhibitor screening. Here we describe the use of 2-AP in an easy-to-use exonuclease assay suitable for high-throughput screening to measure exonuclease activity in real time. Due to its simplicity and minimal use of protein and substrate, this assay is suitable for screening of large chemical libraries to search for novel exonuclease inhibitors. We furthermore show an example where the 2-AP exonuclease assay is used to discover a new inhibitor of the exonuclease domain of the replicative DNA polymerase DnaE1 from *M. tuberculosis*.

**Figure 1.**
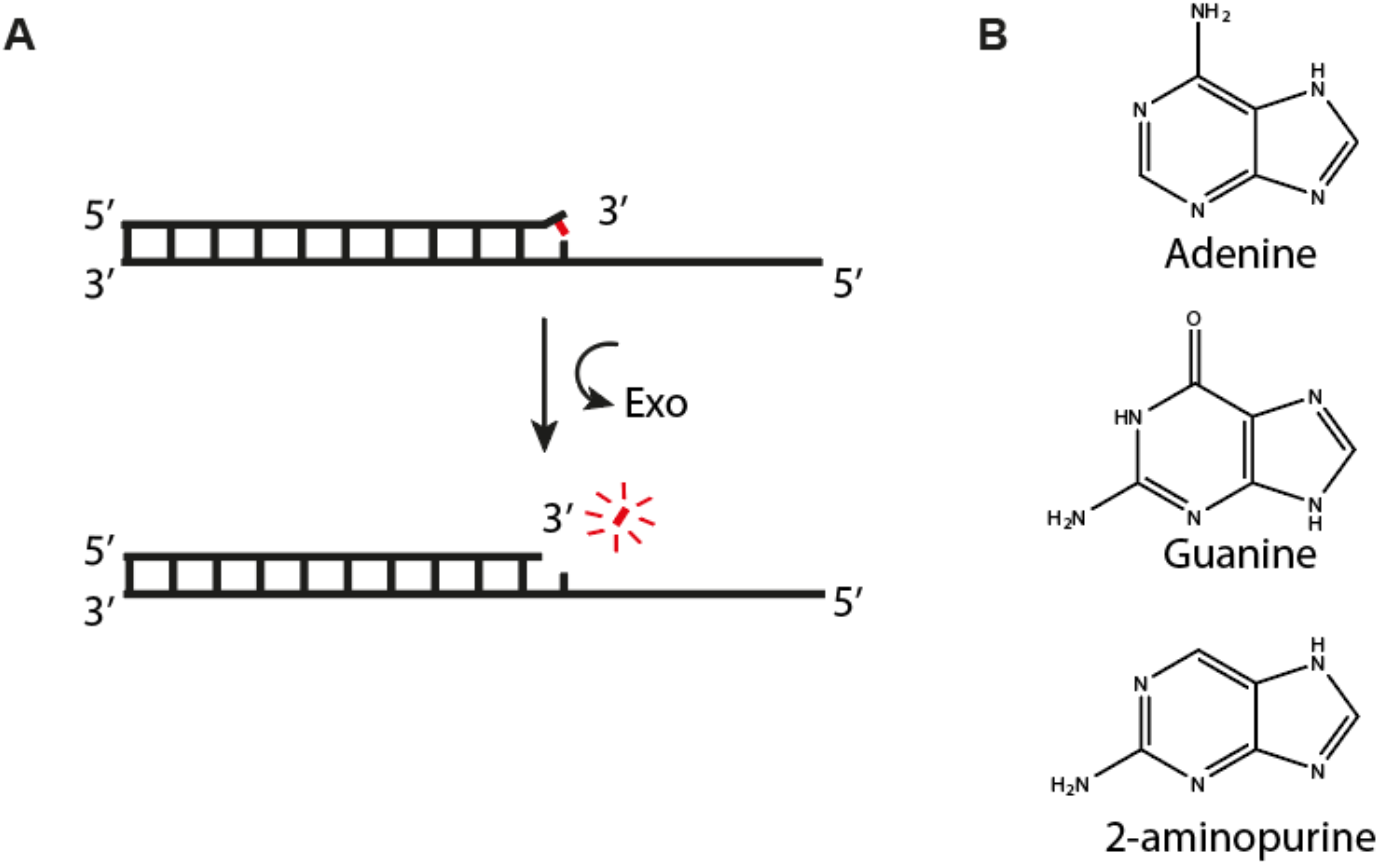
Principle of the 2-aminopurine exonuclease assay. (A) Schematic representation of a DNA substrate containing a 2-AP nucleotide positioned at the 3’ end, where its fluorescence is quenched by the neighbouring bases. Upon cleavage by an exonuclease, the 2-AP is released from the DNA and its fluorescence restored. (B) Comparison of the chemical structures of adenine, guanine, and 2-aminopurine.

## MATERIALS AND METHODS

### Chemicals and reagents

All chemicals were purchased from Sigma, unless indicated otherwise. DNA oligonucleotides were purchased from IDT. Chromatography columns were purchased from Cytiva.

### Site directed mutagenesis

Site direct mutagenesis was used to create the two active site mutants of the *E. coli* exonuclease ε. The Methionine 18 to alanine (ε^18^) was created using the following forward primer 5’-GAAACCACCGGTGCGAACCAGATTGGTGCG-3’ and reverse primer 5’-CGCACCAATCTGGTTCGCACCGGTGGTTTC-3’. The valine 65 to alanine mutation (ε^65^) was created using the following forward 5’-GGAAGCCTTTGGCGCACATGGTATTGCCGATG-3’ and reverse primer 5’-CATCGGCAATACCATGTGCGCCAAAGGCTTCC-3’.

### Protein expression and purification

All proteins were expressed in *E. coli* BL21 (DE3) (Novagen) at 37 °C for 2 hours, unless otherwise stated. Protein purification was performed using buffer A (25 mM Imidazole pH 7.5, 500 mM NaCl, 2 mM DTT, 5% Glycerol), buffer B (25 mM Tris pH 8.5, 2 mM DTT, 10% glycerol), buffer C (25 mM HEPES pH 7.5, 100 mM NaCl, 2 mM DTT), and buffer D (25 mM Tris pH 8.5, 2 mM DTT, 10% glycerol) with addition of imidazole or NaCl as indicated. All purified proteins were flash frozen in liquid nitrogen and stored at -80 °C.

Since *E. coli* ε is expressed in the insoluble fraction, after cell lysis and centrifugation the pellet was solubilized in buffer A with 6 M urea and centrifuged at 24000 x g. The supernatant was injected onto a Histrap column pre-equilibrated in buffer A with 6 M urea and eluted using a gradient to 500 mM Imidazole in buffer A with 6 M urea. The eluted protein was diluted to 0.5 mg/ml in 25 mM Tris, 3 M Urea and 2 mM DTT and refolded by overnight dialysis to buffer B. The refolded protein was centrifuged at 24,000 x g and injected onto a Hitrap Q column pre-equilibrated in buffer B with 40 mM NaCl and eluted with a gradient to 1 M NaCl in buffer B. *E. coli* Pol I and Pol IIIα were purified using a HisTrap column pre-equilibrated in buffer A and eluted using a gradient to 500 mM Imidazole in buffer A. The his-tag was removed by overnight digestion with PreScission Protease (Cytiva), buffer exchanged to buffer A and followed by a second Histrap column to remove undigested protein. The flowthrough was injected onto a Hitrap Q column pre-equilibrated in buffer B with 150 mM NaCl and eluted with a gradient to 1 M NaCl in buffer B. Pol IIIα was further purified using a Superdex 200 size exclusion column pre-equilibrated in buffer C.

*M. tuberculosis* DnaE1 was expressed in *M. smegmatis* cells at 37 °C for 72 hours. It was purified using a HisTrap column pre-equilibrated in buffer A and eluted using a gradient to 500 mM Imidazole in buffer A. The his-tag was removed by overnight digestion with PreScission Protease (Cytiva), buffer exchanged to buffer A and followed by a Heparin column. Lastly it was further purified using a Superdex 200 size exclusion column pre-equilibrated in buffer C.

The catalytic domain of human Exo1 was expressed in *E. coli* BL21 (DE3) overnight at 18 °C ^20^. It was purified using a HisTrap column pre-equilibrated in Buffer C with 20 mM Imidazole and eluted using a gradient to 300 mM Imidazole. Then it was further purified using a Hitrap Q column equilibrated in Buffer C and eluted with a gradient to 1 M NaCl in Buffer C.

### Fluorescence intensity experiments

Assays were performed using oligonucleotides in Table S1. Fluorescence emission data were collected using a PHERAstar FSX microplate reader. All the substrates and the proteins are individually diluted in 50 mM HEPES pH 7.5, 100 mM potassium glutamate, 5 mM MgCl2 and 0.5 mg/ml BSA, unless otherwise stated. Reactions were started by adding protein to DNA in a Corning® 384-well Low Volume Black Round Bottom Polystyrene NBS Microplate (Corning #4514). Data are collected for 50 cycles each lasting 20 seconds. All the steps are performed at room temperature.

The samples were excited at 330 nm and the emission was collected at 380 nm (for excitation and emission spectra, see Figure S1). A 320-380 filter (1904A1 BMG Labtech) was used to collect the measurements. To be able to compare measurements taken on different days, with different exonucleases and on different substrates, all assays were performed using the same gain.

### High throughput screening assays

For the high throughput screening assays, we used the same oligonucleotides used for the manual assays (see Table S1). The oligonucleotides and protein (*M. tuberculosis* DnaE1) were diluted in 50 mM HEPES pH7.5, 100 mM potassium glutamate, and 0.5 mg/mL BSA. Assays were dispensed using a Mantis liquid handler (Formulatrix) in 384 well polypropylene plates from Corning # 4514. Data collected for 30 cycles each lasting for 60 seconds.

The assay validation statistics (Figure S5) were calculated using the following formulas:

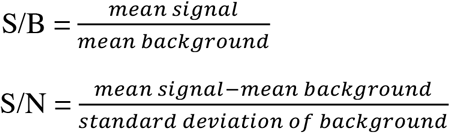

Where the mean signal is the fluorescent signal at completion of the assay in presence of DNA and protein. The mean background is the negative control, where only DNA was added.

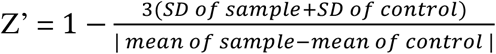

Where ‘SD of sample’ is the standard deviation of the positive control (DNA + protein) and the ‘SD of control’ is the standard deviation of the negative control (DNA only).

### Differential Scanning Fluorometry assays

The melting temperature of the *M. tuberculosis* DnaE1 was determined in the absence and presence of the ligand 6-thioinosine using a Tycho instrument (Nanotemper). 8 *μ*M of *M. tuberculosis* DnaE1 was incubated with 100 *μ*M 6-thioinosine for 10 minutes at 20 °C. Fluorescence intensity was followed at 330 and 350 nm while the temperature was increased from 35 to 95 °C over 3 minutes. Data was analyzed with Tycho software.

## RESULTS

### Optimization of assay conditions

In order to define a suitable working range for the 2-AP exonuclease assay, we analyzed the assay under different DNA, protein, and salt concentrations, as well as different pH values. For this we use the *E. coli* exonuclease ε, a well characterized enzyme responsible for removal of mis-incorporated nucleotides during DNA replication^21–24^. Because the exonuclease ε works in a 3’ to 5’ direction, experiments were performed using a substrate containing a mismatched 2-AP at the 3’ end (substrate 2AP11 in Table S1). Using a range of DNA concentrations, from 30 nM to 20 *μ*M and a fixed amount of protein (0.5 nM), we find a clear signal over the entire range, indicating the assay is sensitive over for a wide range of substrate concentrations (Figure 2A). Therefore, to limit the amount of DNA consumed, we used 0.5 *μ*M DNA in all subsequent assays. Next, we determined the enzyme working range by varying the concentration of ε from 0.125 -4 nM (Figure 2B). Here too we find that the assay gives a robust signal, even at concentrations as low as 0.125 nM. In addition, as salt can influence the interaction between many enzymes and DNA, we also varied the salt concentration while keeping the concentration of DNA and ε fixed at 0.5 *μ*M and 0.5 nM, respectively (Figure 2C). Here we find that for ε there is an optimal salt concentration of ∼50 mM potassium glutamate, whereas no activity was observed at low (0 mM) or high (500 mM) salt concentrations. However, because the observed salt concentration for a bacterial cell is 100-200 mM, we decided to maintain a salt concentration of 100 mM for all subsequent experiments. Finally, as it has been reported that pH influences the fluorescence intensity of 2-AP^25^, we also measured the difference between free 2-AP and 2-AP incorporated into a DNA substrate over a wide range of pH values, from pH 2.5 to pH 10 (Figure 2D, and Table S2). We find that the 8-to 9-fold difference between free 2-AP and DNA-bound remains stable between pH 5 and pH 10 and only drops to a 4-fold difference at pH 2.5. Hence the assay is also applicable over a wide range of pH values.

**Figure 2.**
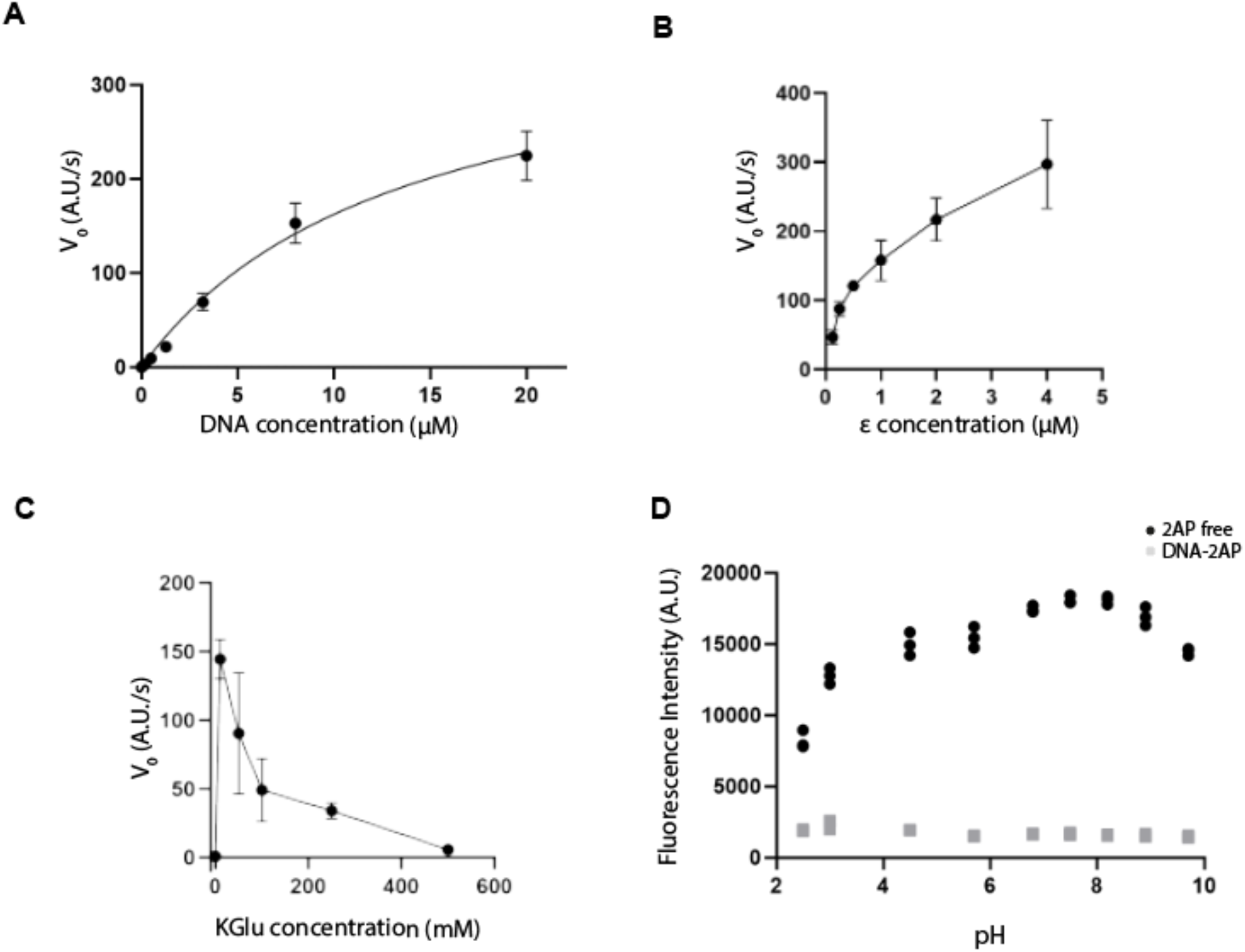
Determination of suitable working range for the 2-AP exonuclease assay. (A) Initial velocities (V_0_) for the 2-AP exonuclease assay at 0.03 - 20 *μ*M DNA and 0.5 nM *E. coli* ε exonuclease. Raw data shown in Figure S2 (B) Initial velocities (V_0_) for the 2-AP exonuclease assay performed using 0.125 - 4 nM *E. coli* ε exonuclease and 0.5 *μ*M DNA. (C) 2-AP exonuclease assay at 0 - 500 mM potassium glutamate (KGlu). Experiments performed using 0.5 *μ*M DNA and 0.5 nM ε. (D) Difference between free 2-AP (black circles) and DNA-bound 2-AP (grey squares) over a pH range from 2.5 - 10. Buffers used are listed in Table S2. All data are derived from three or more independent experiments. Raw data is shown in Figure S2 and S3.

### Versatility of the assay across different exonucleases families

To further test the usefulness of the 2-AP exonuclease assay, we measured the activity two *E. coli* ε mutants, ε^18^ and ε^65^, alongside that of wild type ε (Figure 3A-B). Both exonuclease mutants contain a mutation positioned at the entrance of the active site, and have been shown to reduce exonuclease activity ^26^. As anticipated, the two mutants ε^18^ and ε^65^ show a reduction in activity by 80% and 66%, respectively, indicating that the 2-AP assay is useful for characterization of different mutant version of a protein.

**Figure 3.**
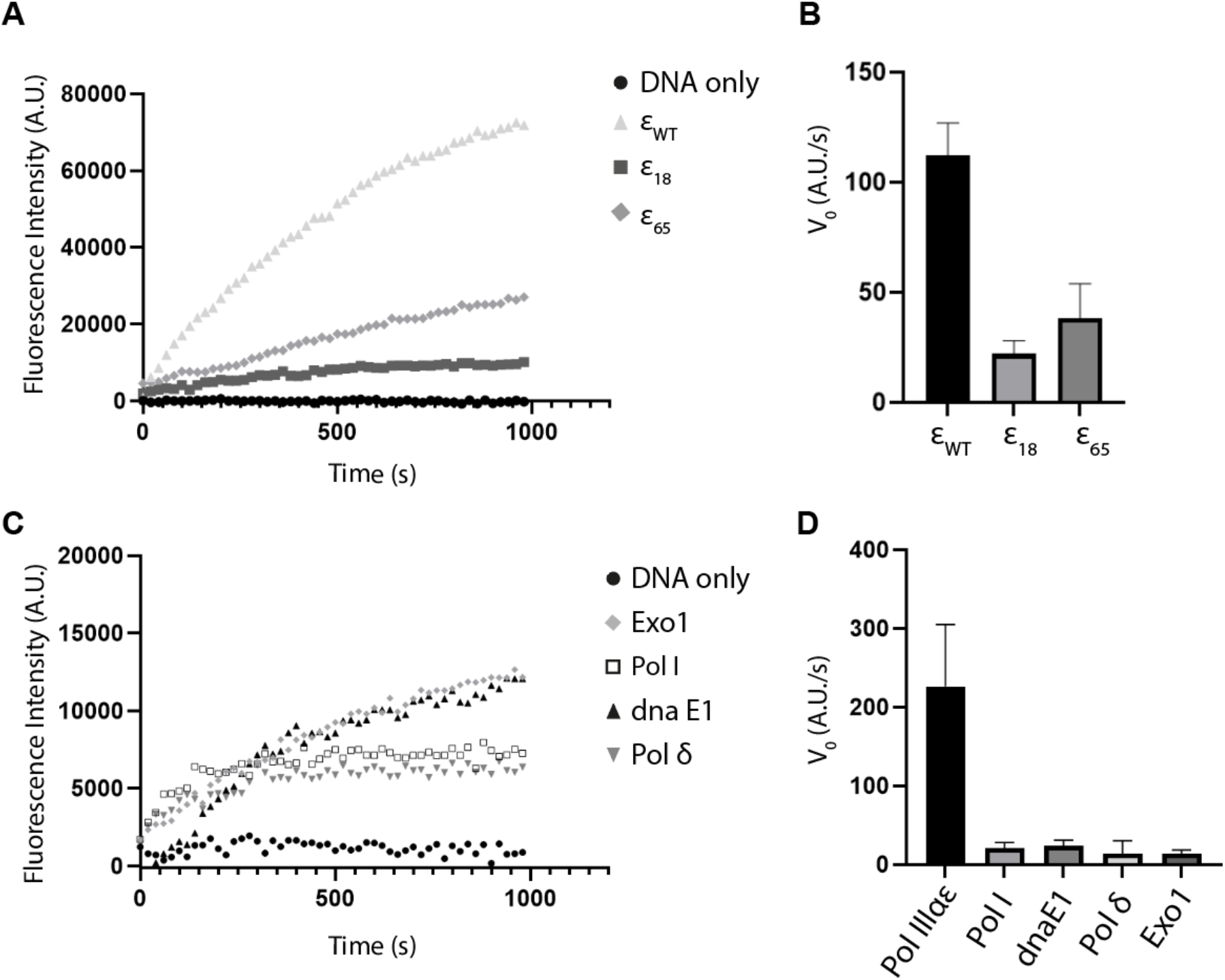
2-AP exonuclease assay used on a variety of different proteins. (A) 2-AP exonuclease activity of wild type ε and two active site mutants, ε^18^ and ε^65^. Assays performed using 0.5 nM ε and 500 nM 2AP11 substrate. (B) Bar graph showing the initial velocity (V_0_) for the curves shown in A. (C) 2-AP exonuclease assay performed with *E. coli* Pol I, *M. tuberculosis* DnaE1, human Pol δ and human Exo1. All proteins were used at 0.5 nM, except for Exo1 that was used at 10 μM. 500 nM of substrate was used for all the proteins. For Exo1 that is a 5’-3’ exonuclease a DNA substrate with a 2-AP positioned at the 5’ was used (2AP11 Reverse in Table S1). (D) Bar graph showing the initial velocity (V_0_) for the curves shown in panel C. For comparison, the V_0_ of *E. coli* ε is included.

To further explore the versatility of the assay we tested different 3’-5’ exonucleases from different organisms as well as a 5’-3’ exonuclease. For this we used *E. coli* Pol I, *M. tuberculosis* DnaE1, human Pol δ and the human 5’-3’ exonuclease Exo1 (Figure 3C). Remarkably, these exonucleases show a reduced activity, between 6 - 11% of that of *E. coli* ε (Figure 3D). This is, however, in agreement with previous literature ^2,27–29^.

In the assay conditions used above (0.5 nM protein, 0.5 *μ*M DNA), most curves exit the linear phase at ∼10 minutes, which is a reaction time that is too short when screening large compound libraries where several automated steps need to be executed sequentially for a large number of plates. Therefore, to slow down the reaction we searched for an alternative way to influence the reaction velocity, as the protein concentration was already low (0.5 nM) and the DNA concentration a 1000-fold higher (500 nM). Previously, using a gel-based exonuclease assay we observed that the excision of a 3’ terminal 2-AP nucleotide was faster than a regular nucleotide (data now shown). This faster excision of the 2-AP may be problematic when searching for potential exonuclease inhibitors as weak inhibitors of ‘normal’ exonuclease activity may not be effective on the faster 2-AP, and therefore will not be picked up during the screening of a chemical library. We therefore wondered if placing the 2-AP further away from the 3’ end could slow down the reaction as the exonuclease would first have to remove regular nucleotides before excising the 2-AP (Figure 4A). Indeed, moving the 2-AP two or three nucleotides away from the 3’ end resulted in a ∼10 to 60-fold decrease of the exonuclease activity, respectively (Figure 4A-B). We therefore choose the DNA substrate with the 2-AP positioned two nucleotide away from the 3’ end (i.e. 2AP14) for all further experiments.

**Figure 4.**
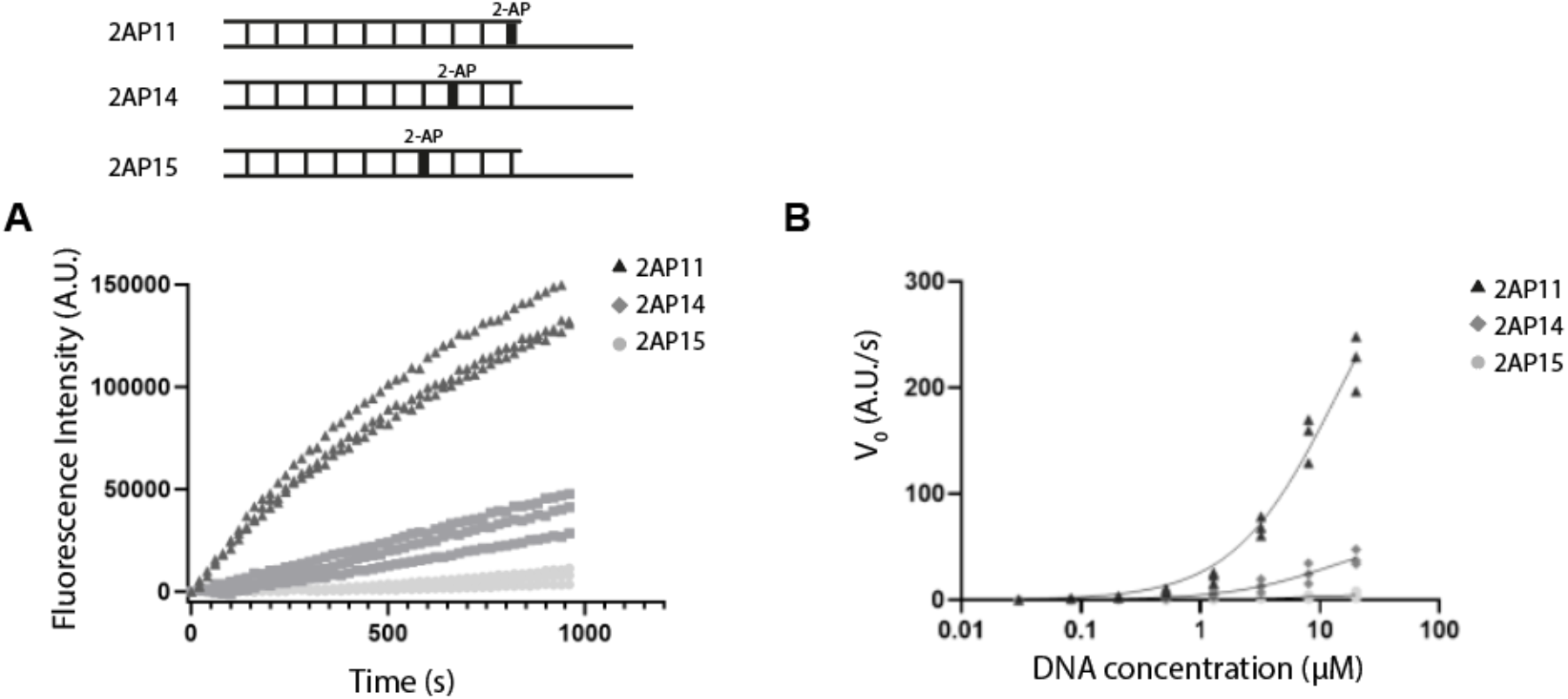
Position of 2-AP affects exonuclease activity. (A) Comparison of *E. coli* ε exonuclease activity using DNA substrates with the 2-AP at three different positions (highlighted in bold). Assay performed using 20 μM of each substrate and 0.5 nM ε exonuclease. Data from three independent experiments (B) Graph showing initial slope velocity (V_0_) of reactions performed using 1 nM ε and increasing DNA concentration (0.08 to 20 μM). Data from three independent experiments.

### High throughput inhibitor screening adaptability

Finally, to test the suitability of the 2-AP exonuclease as a high-throughput inhibitor screening tool we chose the PHP-exonuclease domain of the replicate DNA polymerase DnaE1 from *M. tuberculosis* as a target for our inhibitor screen. The PHP-exonuclease domain of DnaE1 is an attractive target for novel antibiotics as it is biochemically and structurally distinct from the human exonucleases^30^ and essential for viability in *M. tuberculosis*^31^. As the exonuclease activity of *M. tuberculosis* DnaE1 is lower than that of *E. coli* ε (Figure 3D), we increased the protein concentration from 0.5 nM to 15 nM to create an assay speed that is suitable for our high throughput inhibitor screen setup. In addition, we used a Mantis (Formulatrix) automated liquid handling robot for rapid dispensing and reduction of the reaction volume from 15 to 10 μ L. Under these conditions, we find that the mean fluorescence intensity signal in presence of enzyme is 243.8 × 10^3^ ± 6.7 × 10^3^ while the background fluorescence intensity signal in absence of enzyme is 78.2 × 10^3^ ± 6.3 × 10^3^, a three-fold difference between the signal and background. This yields a Z-score of 0.8, which, together with other quality indicators (Figure S5), indicates a highly reliable confidence level^32^ for the 2-AP exonuclease assay. In addition, we also tested a positive control for inhibition of exonuclease activity. For this we used a DNA substrate that contains a non-hydrolysable phosphorothioate bond between the 3’ terminal nucleotide and adjacent nucleotide, which acts as a competitive inhibitor to the 2-AP substrate (Figure 5A and substrate ssDNA-P in Table S1). Already at a concentration that is ten-fold less than the substrate DNA, the phosphorothioate DNA reduces the exonuclease activity by ∼30% indicating that it is a suitable positive control for exonuclease inhibition.

**Figure 5.**
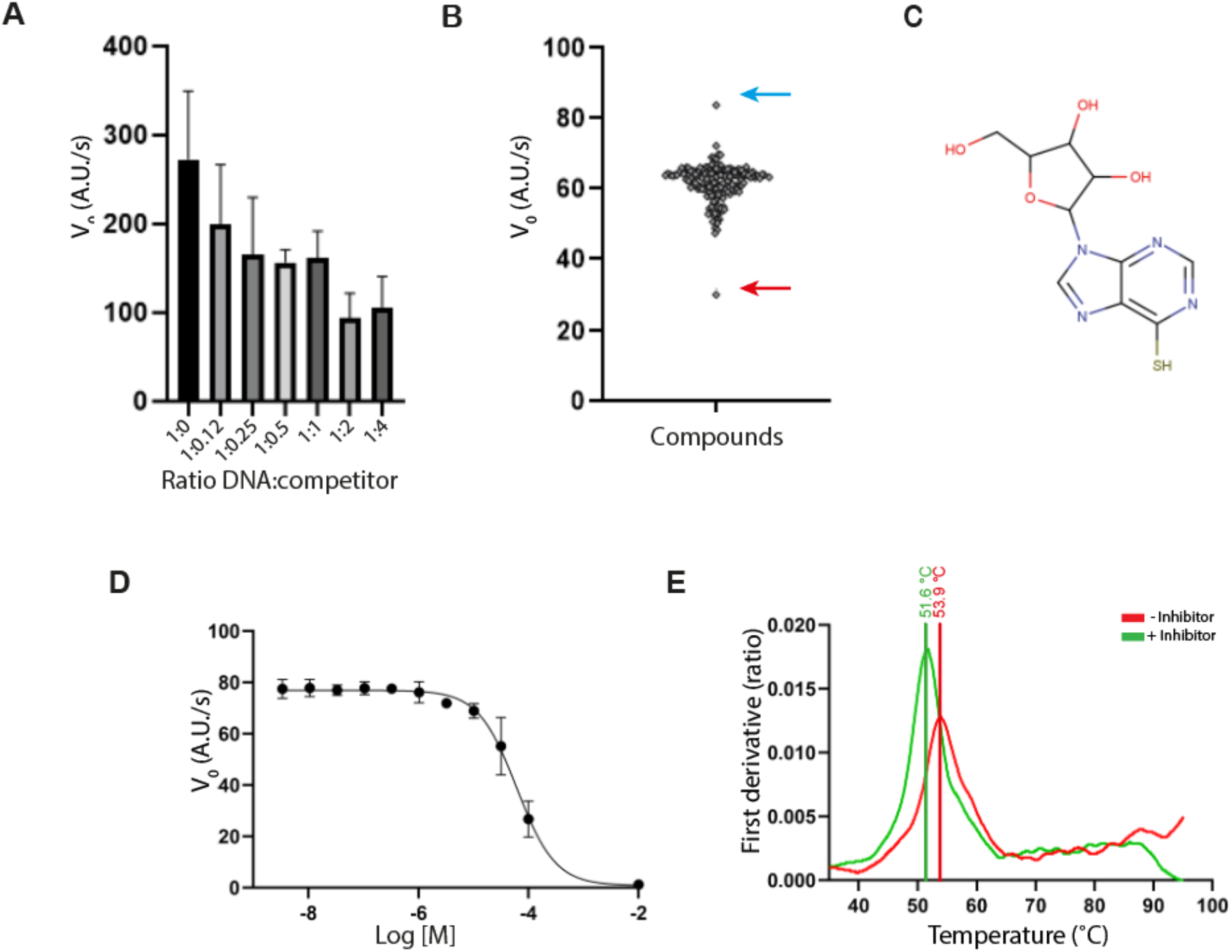
High throughput screening using the 2-AP exonuclease assay (A) Initial slope velocities (V_0_) for a 2-AP assay performed using 15 nM *M. tuberculosis* DnaE1 and 1 μM of 2AP14 substrate and 0.12 - 4 μM of inhibitor DNA. (B) Violin plot showing the initial velocity values for each assay reaction in the respective wells. Values of V_0_ around 65 indicate that there is no inhibition of protein activity. The red arrow indicates the hit compound 6-thioinosine. The blue arrow indicates a compound that shows intrinsic fluorescence. (C) Chemical structure of 6-thioinosine (D) Dose response curve of 6-thioinosine’s inhibitory action on *M. tuberculosis* DnaE1 exonuclease activity. (E) First derivate plot of a thermal melt measurement of *M. tuberculosis* DnaE1 in absence (red line) and presence (green line) of 6-thioinosine. Raw data in Figure S4B.

Finally, to screen for potential inhibitors of the *M. tuberculosis* DnaE1 exonuclease we used a nucleotide compound library from MedChemExpress that contains ∼200 nucleotide and nucleoside analogues. Using this library, we were able to identify a single hit compound, 6-thioinosine, which inhibits the exonuclease activity of *M. tuberculosis* DnaE1 by ∼50 % at 100 μM concentration (Figure 5B-C). The compound was further characterized by a dose-response curve (Figure 5D) yielding an IC_50_ of 64 μM. Binding of the 6-thioinosine to DnaE1 was also validated using differential scanning fluorometry that revealed a two-degree shift in the melting temperature of the protein (Figure 5E, Figure S4B). While the affinity of 64 μM is too low for a suitable drug candidate, it provides an attractive starting point for development of better inhibitors against the exonuclease site of DnaE1 from *M. tuberculosis*. Finally, it is worth mentioning that some compounds show intrinsic fluorescence and may obscure the signal (Figure 5B). It is therefore helpful to also screen the compound library in absence of enzyme to identify any intrinsically fluorescent candidates that need to be excluded from the results.

### Discussion and concluding remarks

As the threat of drug-resistant and multi-drug resistant bacteria is increasing world-wide^33^ the need for novel antibiotics with new mode of actions is higher than ever. While inhibition of genome replication is frequently used in antiviral therapies, there are currently no antibiotics that directly target DNA replication in bacteria. Several inhibitors against bacterial replication proteins have been described, such as the *E. coli* DNA helicase DnaB, the sliding clamp β, the RNA primase DnaG and the replicative DNA polymerases Pol C and DnaE^34–36^, yet there are currently none that have been developed into an antibiotic that is approved for use by the general public. Therefore, the need for novel DNA replication inhibitors remains. Replicative exonucleases are attractive targets for novel antibiotics, as disruption of exonuclease activity dramatically compromises viability in *E. coli, S. typhimurium*, and *M. tuberculosis*^23,24,31^. Moreover, the majority of bacteria employ a so-called PHP exonuclease^31^ that is only found in bacteria and has no known homologs in eukaryotes^37^, thus reducing the likelihood of cross-species inhibition.

Exonucleases are also of interest as potential targets of anticancer drugs. More than ten exo- and endonucleases are active in different forms of mammalian DNA repair (reviewed in^38^). Furthermore, upregulation of EXO1 is associated with increased malignancies and correlated with a poor prognosis ^39,40^, whereas the inhibition of APE2 has shown to reduce growth and malignancies ^41,42^. Finally, also ExoN, the replicative exonuclease of the SARS-CoV-2 virus is the target of novel promising antiviral drugs ^43^.

To aid in the search of novel exonuclease inhibitors we have developed a simple and versatile exonuclease assay that can be used for both 3’-5’ and 5’-3’ exonucleases, as well as bacterial and human exonucleases. Using this assay that is compatible with high-throughput inhibitor screening we have discovered a novel inhibitor of the *M. tuberculosis* DnaE1 exonuclease that is essential for viability in this bacterium. Hence our work describes a novel, real-time exonuclease assay that can be used to search for novel inhibitors of DNA replication of DNA repair.

## Acknowledgments

Pol δ was a kind gift from the laboratory of Prof. Hamdan, at King Abdullah University of Science and Technology, Saudi Arabia. The manuscript was with written with contributions of all authors. All authors have given approval to the final version of the manuscript.

## Notes

### Competing Interest Statement

The authors have declared no competing interest.

