## Supplementary Material for "A high-throughput exonuclease assay based on the fluorescent base analog 2-aminopurine"

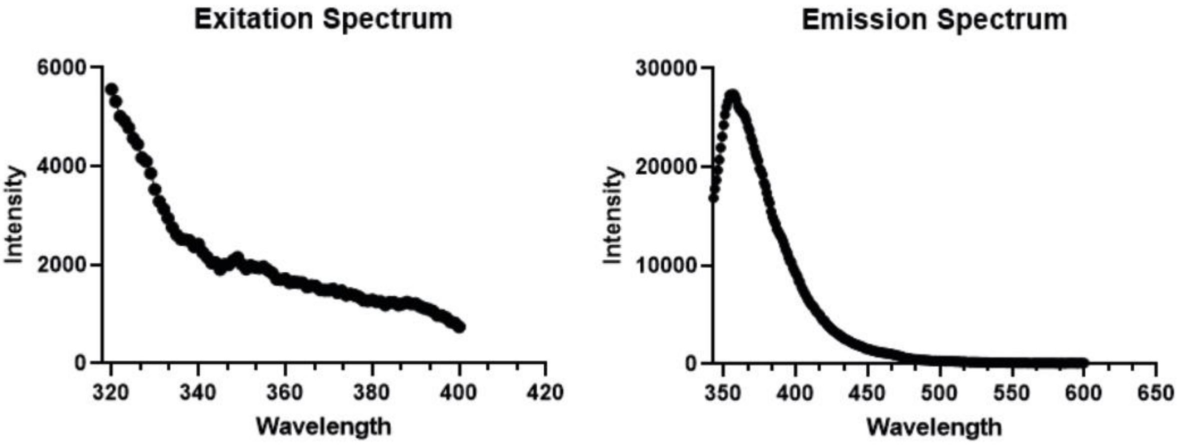


**Figure S1.** Excitation and Emission spectrum of 2-AP

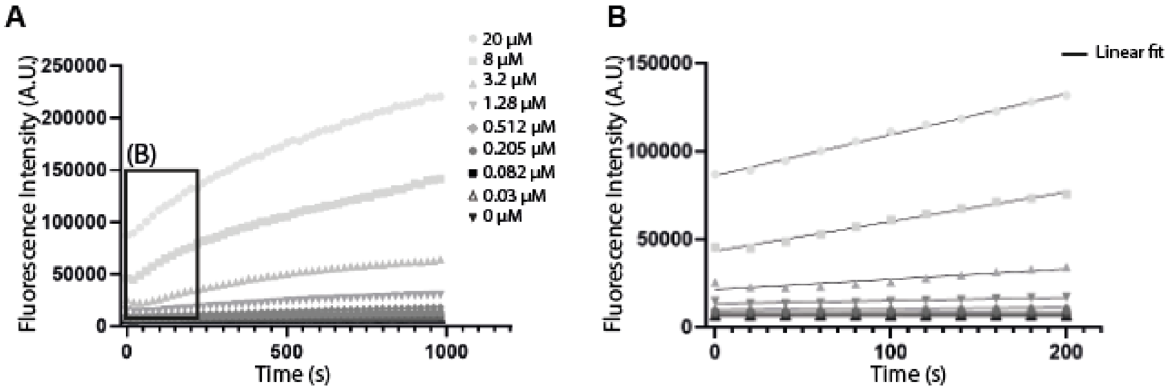


**Figure S2.** (A) 2-AP exonuclease assay performed over a titration of DNA. (B) Close up of the first 200 seconds marked by a square in in panel A. The linear section of the curve was used to measure the initial velocities, $V_0$.

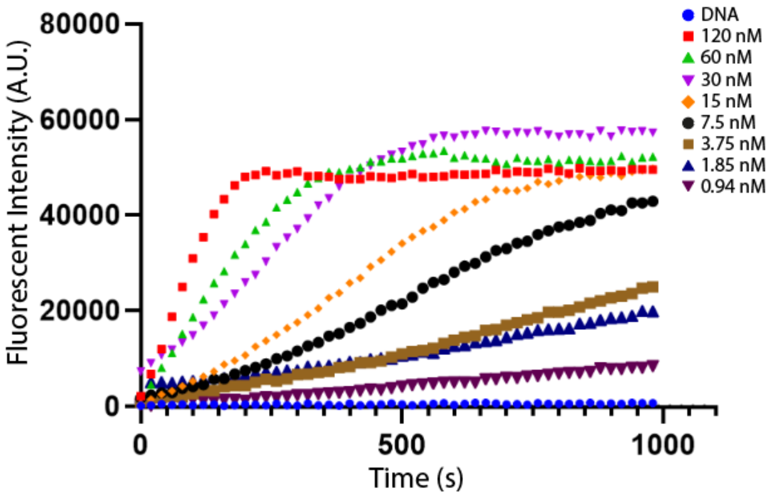


**Figure S3.** 2-AP exonuclease assay performed over a titration of DnaE1 from 0.9 nM to 120 nM. 500 nM of 2AP14 DNA was used.

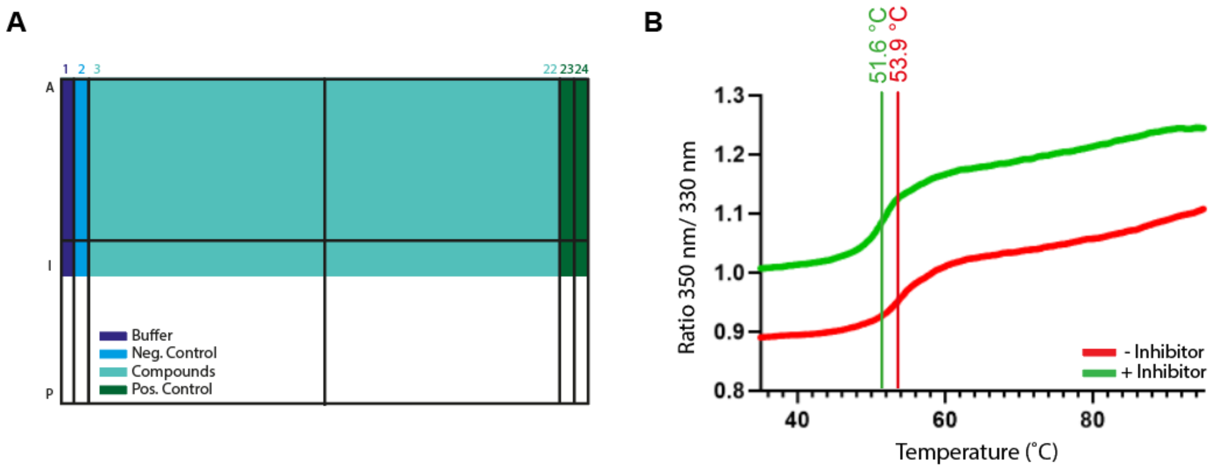


**Figure S4.** (A) Representation of the layout used for high throughput screening assays. The plate contains, in addition to the ~200 nucleotide analogs, also a negative (i.e. DNA alone) and a positive control (the non-hydrolysable phosphorothioate DNA) (B) Differential scanning fluorometry measurements of DnaE1 alone (red curve) and DnaE1 in presence of the inhibitor 6-thioinosine (green curve). The presence of 6-thioinosine results in a decrease of the melting temperature by two degrees.

A

| | Plate 1 | Plate 2 | Plate 3 |
|---|---|---|---|
| Z' | 0,91269 | 0,87080 | 0,83039 |
| S/B | 3,51542 | 3,21548 | 3,34554 |
| S/N | 1199,94 | 295,317 | 80,4496 |

B

| | Day 1 | Day 2 | Day 3 |
|---|---|---|---|
| Z' | 0,90089 | 0,67939 | 0,83039 |
| S/B | 3,03119 | 3,27630 | 3,34554 |
| S/N | 72,438 | 64,697 | 80,4496 |

**Figure S5.** Assay validation. (A) Values for the Z', S/B and S/N calculated over three independent data sets acquired on the same day. (B) Values for the Z, S/B and S/N calculated over three independent data sets acquired on three different days.

| **Name** | **Sequences** |
|---|---|
| 2AP11 | 5'GTTCACGAGACCTACTGACACTG**A** 3'<br>3'CAAGTGCTCTGGATGACTGTGACGTGTACGTA 5' |
| 2AP11 Reverse | 5'**A**TTCACGAGACCTACTGACACTGC 3'<br>3'CAAGTGCTCTGGATGACTGTGACGTGTACGTA 5' |
| 2AP14 | 5'GTTCACGAGACCTACTGACACT**A**AG 3'<br>3'CAAGTGCTCTGGATGACTGTGAGTCGTACGTA 5' |
| 2AP15 | 5'GTTCACGAGACCTACTGACAC**A**GAG 3'<br>3'CAAGTGCTCTGGATGACTGTGGCTCGTACGTA 5' |
| ssDNA-P | 5'GGAGTAGTACTAGGACGAAGGACTC*T 3' |

**Table S1** Primers used for fluorescence intensity experiments. The position of 2-aminopurine nucleotide analog is highlighted in bold. Asterisk points the position of the non-hydrolysable phosphorothioate bond.

| Buffer | pH |
|---|---|
| Citric Acid | 2.5 |
| Citric Acid | 3 |
| Citric Acid | 4.5 |
| MES | 5.7 |
| HEPES | 6.8 |
| HEPES | 7.5 |
| HEPES | 8.2 |
| HEPES | 8.9 |
| Glycine | 9.7 |

**Table S2.** Buffers used for assay in Figure 2D. Final concentration of the buffer in the assays was 50 mM.
